# Model-checking ecological state-transition graphs

**DOI:** 10.1101/2021.11.19.469210

**Authors:** Colin Thomas, Maximilien Cosme, Cédric Gaucherel, Franck Pommereau

## Abstract

Model-checking is a methodology developed in computer science to automatically assess the dynamics of discrete systems, by checking if a system modelled as a state-transition graph satisfies a dynamical property written as a temporal logic formula. The dynamics of ecosystems have been drawn as state-transition graphs for more than a century, from state-and-transition models to assembly graphs. Thus, model-checking can provide insights into both empirical data and theoretical models, as long as they sum up into state-transition graphs. While model-checking proved to be a valuable tool in systems biology, it remains largely underused in ecology. Here we promote the adoption of the model-checking toolbox in ecology through its application to an illustrative example. We assessed the dynamics of a vegetation model inspired from state-and-transition models by model-checking Computation Tree Logic formulas built from a proposed catalogue of patterns. Model-checking encompasses a wide range of concepts and available software, mentioned in discussion, thus its implementation can be fitted to the specific features of the described system. In addition to the automated analysis of ecological state-transition graphs, we believe that defining ecological concepts with temporal logics could help clarifying and comparing them.

**Author summary:** Ecologists have drawn state-transition graphs representing the dynamics of ecosystems for more than a century. Model-checking is an automated method for the analysis of such graphs developed in computer science and acknowledged by a Turing award in 2007. Ecologists appear to be mostly unaware of model-checking despite its successes in systems biology to assess the dynamics of biological networks.

We promote model-checking of ecological state-transition graphs through its application to an illustrative vegetation model. We exemplify the insights provided by model-checking by assessing management policies aiming to tackle savanna encroachment. We also provide a catalogue of patterns to help ecologists with the difficulty of formally expressing dynamical properties. We also discuss the wide range of model-checking concepts and available software, enabling to fit the specific features of the studied system, such as durations or probabilities.

Model-checking can be applied to both empirical data and theoretical models, as long as they sum up into state-transition graphs. It provides automated and accurate answers to complex questions that could barely be analysed through human examination, if not impossible to answer this way. In addition to the automated analysis of ecological state-transition graphs, we believe that formally defining ecological concepts within the model-checking framework could help in clarifying and comparing them.

## Introduction

A *state-transition graph (STG)* describes the behaviour of a dynamical system as discrete states linked by transitions. Ecologists have drawn STGs for more than a century, one of the earliest and best known example being the vegetation successions described by Clements [1]. Yet, ecology and environmental sciences appear to remain largely unaware of the *model-checking* methodology [2] developed by computer scientists to investigate the dynamics of a system represented as a STG. This paper aims to promote the model-checking of ecological STGs through its application to an illustrative example taken from the literature [3].

In ecology, STGs are typically used to represent the *community pathways*, i.e. the changes in the set of species or populations, of an ecosystem through time. Such as for example the successions of plant communities in boreal forests [4], or the assembly of microbial communities in laboratory experiments [5]. Such STGs are mostly drawn from observations, hence their relatively small size (a few dozens of states at most). Most of the time, STGs are perceived as graphical representations of the knowledge about the dynamics of the studied system rather than as actual data.

In addition, STGs have been used since the 90’s as a tool for rangeland management and ecosystem conservation under the concept of *state-and-transition models (STMs)* [6, 7]. Theoretical studies also emphasise the relevance of STGs to investigate community assembly [8, 9]. Both research fields recently mentioned an interest in tools providing dynamical analysis of STGs [9, 10].

In computer science, STGs model the execution of automated systems. Computer scientists design automated tools called *model-checkers* to ensure the absence of bugs during software executions [2]. Model-checkers verify whether the pathways within the STG model satisfy a given property, for example that a desired state remains reachable or that a harmful behaviour is always avoided. Model-checking is an active field of research acknowledged by a Turing award in 2007, encompassing numerous concepts and resulting in a wide variety of implemented tools [11, 12].

In systems biology, STGs are usually the outputs of models of reaction networks or regulatory networks [13]. Model-checking is extensively used to analyse those models [14, 15], proving its suitability for the study of biological systems. For example, model-checking helped validating models of nutritional stress response of *Escherichia coli* [16], T-helper cell reprogramming [17], mammalian cell cycle [18] or BRAF inhibition pathways in two different cancers [19].

Yet, model-checking methodology appears to remain unknown to most ecologists despite the need for tools analysing STGs. Indeed, ecology encompasses a wide variety of STGs, from state-and-transition models to assembly graphs, but their analysis is often restricted to visual examination. Meanwhile systems biology proved that model-checking can provide valuable insights to biological systems. This article promotes the model-checking of ecological STGs, through an illustrative implementation on an example taken from the STM literature [3]. The model-checking methodology can be implemented in a wide variety of ways, that will be covered in Discussion below.

## 1 Materials and methods

### 1.1 State-transition graphs (STGs)

A *state-transition graph (STG)* represents the dynamics of a system as a *graph*, i.e. a set of *nodes* (the discrete states of the system) and *edges* (the transitions enabling to move from one state to another). In ecology, the state of an ecosystem is often discretely abstracted by its *community* (i.e. restricted to its set of species or populations). Subsequently STGs are found in a broad variety of studies focusing on the dynamics of ecological communities, historically called community succession for plants and community assembly for animals [20, 21].

Graphs are widespread in ecology, but STGs must be discriminated from *interaction networks* such as the iconic trophic networks [22, 23]. Indeed, the former grasps the temporal dynamics of an ecological system, while the latter grasps the processes taking place between its components. A node (resp. an edge) of a STG is a temporal stage (resp. an event) of the system dynamics, whereas a node (resp. an edge) of an interaction network is a component (resp. a flux) of the system. The methods presented in this paper deal with the temporal changes of a system and thus are designed to analyse STGs specifically.

#### 1.1.1 State-transition graphs in ecology

The dynamics of ecological systems have been described as states and transitions for more than a century. For example Clements [1] used STGs to represent ecological successions between vegetation states called “seral stages”. Since then, STGs have regularly been used under various names to describe ecological dynamics, from “behaviour graphs” [24], to “kinematic graphs” [25], or under the generic term “pattern” [26].

Moreover, STGs are the core of one of the most commonly used framework for ecological successions: the *state-and-transition models (STMs)*. STMs are derived from observations and have been designed [6] to cope with the non-deterministic and irreversible nature of observed dynamics. STMs have also been designed to be user-friendly, enabling participatory model development and collaborative management [7]. The main goal of STMs is to assist managers and scientists in collectively proposing policies driving the ecosystem through some wanted pathways while avoiding unwanted others. In order to remain user-friendly, STMs sizes usually do not exceed a few dozens of states. While STMs originally stem from rangeland management [6], they are now used in many fields such as natural park management [27] (EDIT STMs database), geomorphology [28], or agroecology [29].

STGs are also found in the field of community assembly under the concept of *assembly graphs*. In such graphs, every node is a stable species community and every transition is an invasion event. Contrarily to STMs, most studies involving assembly graphs are theoretical [8, 9], yet a few are experimental [5].

STGs can also be the output of a wide diversity of modelling formalisms in ecology and environmental sciences [9, 30, 31]. Recently, some studies have used Boolean models such as synchronous *Boolean networks* (i.e. systems of logical equations) to model ecological dynamics, from plant-pollinator associations [32] to spruce budworm outbreaks [33].

#### 1.1.2 Examples from state-and-transition models of the Borana vegetation

The STMs developed by Liao and Clark concern the vegetation pathways of the Borana Zone in southern Ethiopia (Fig. 1). Open canopy woodland (a savanna-like vegetation class encompassing a grass layer with sparse trees) was formerly the most prominent vegetation class in Borana [34]. But since the fire ban in the 1970s the region have been undergoing a rapid increase in the density of woody plants (known as *bush encroachment*). As local people predominantly practice pastoralism, the reduction of herbaceous cover threatens their livelihood. Hence understanding the vegetation pathways is critical for helping pastoralists and policy makers to mitigate bush encroachment [35].

**Fig 1.**
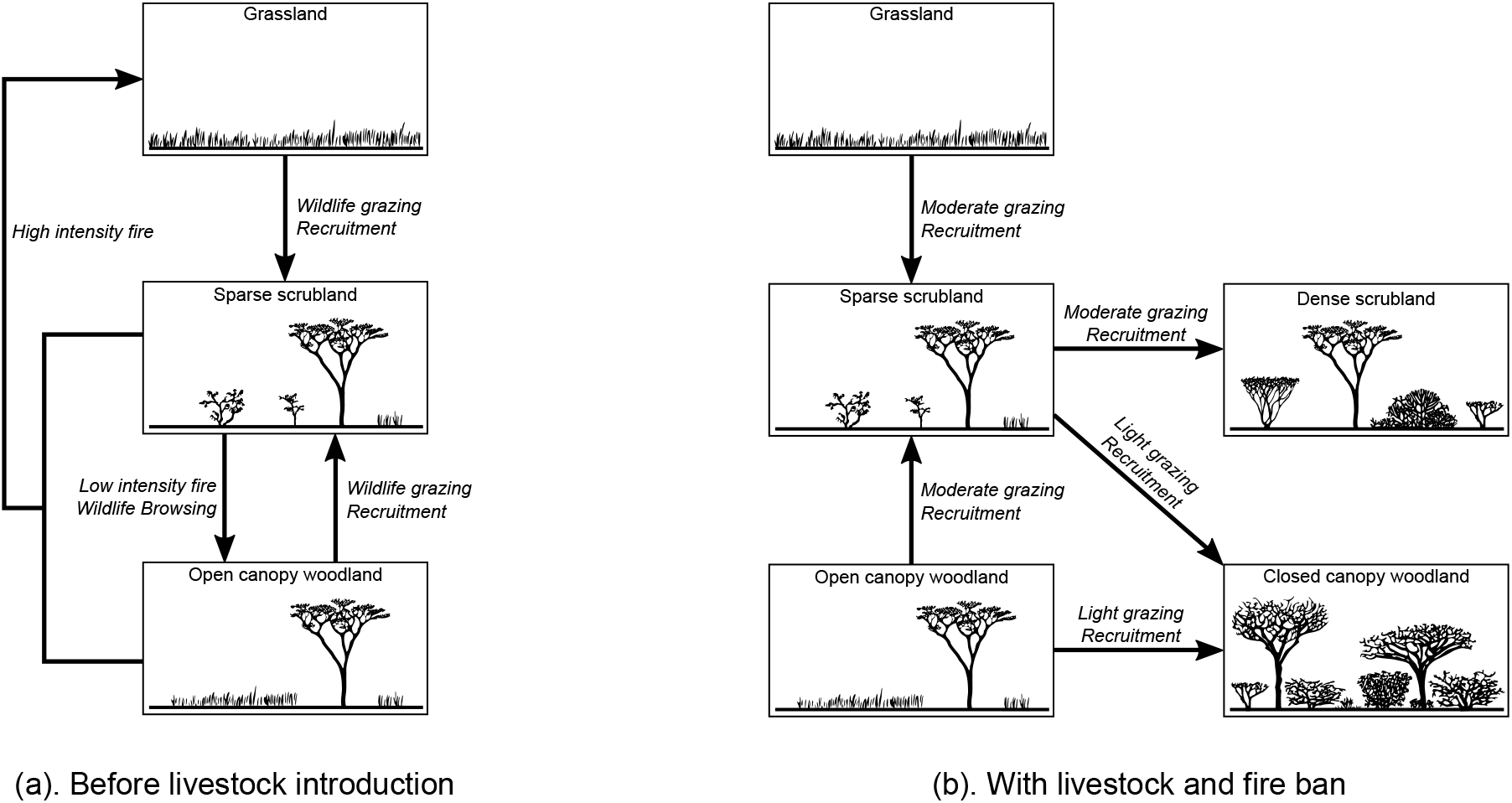
State-and-transition models of the vegetation of the Borana zone. The states are embodied by illustrated boxes, and the transitions by arrows labelled with their main driving processes. (a) Before pastoralism, fire was the main driver of the rangeland dynamics. The combination of fire, wildlife herbivory and vegetation recruitment maintained the entire system in a loop between open canopy woodland and grassland. (b) The presence of cattle and the fire ban gave competitive advantage to woody plants, thus inducing an irreversible bush encroachment. Concurrently, wildlife increasingly avoided the Borana zone because of the denser human and livestock populations. (Based on [3] with author’s permission.)

The states of the STMs represent vegetation classes (Fig. 1), see [34] for their exhaustive definitions. The transitions are labelled with their main drivers as it is often the case in the STM framework. As these STMs consists of nodes and edges representing the vegetation dynamics, they indeed form STGs. The STGs of Fig. 1 are called *non-deterministic* because some states have more than one outgoing transition. Moreover, the STG representing bush encroachment (Fig. 1b) is called *irreversible* because some pathways are one-way only: for example grasslands cannot be reached from any encroached state (dense scrubland or closed canopy woodland).

#### 1.1.3 Building a larger Borana vegetation model with if-then rules

The sizes of the Borana STMs are very limited (Fig. 1). In order to illustrate the scalability of the model-checking methodology, we took inspiration from the literature [3, 34–36] to build a larger model of the Borana ecological successions, called *larger Borana vegetation model* (*LBV-model* for short) in the following.

Each state of the LBV-model consists of a vector of *Boolean variables* representing the functional presence (noted +) or absence (−) of a component of the system. A variable is considered functionally present if its presence has an observable influence on the system, and functionally absent otherwise. Variables influencing the system without being influenced in turn are called *controls*, e.g. climatic conditions or management policies. The transitions are generated from the execution of a rule-based formalism. More precisely, we use *if-then rules* (if the condition is fulfilled, then the consequence may arise), a methodology previously proposed [37, 38] and implemented [39] to model expert knowledge about ecosystem dynamics. Starting from a given state, the outgoing transitions result from the rules whose conditions are fulfilled in this state. Each transition then leads to the state resulting from the application of its consequence to the starting state. This modelling approach is illustrated by a toy model replicating the STM without encroachment (Fig. 1a) and involving only three variables and four rules (Fig. 2).

**Fig 2.**
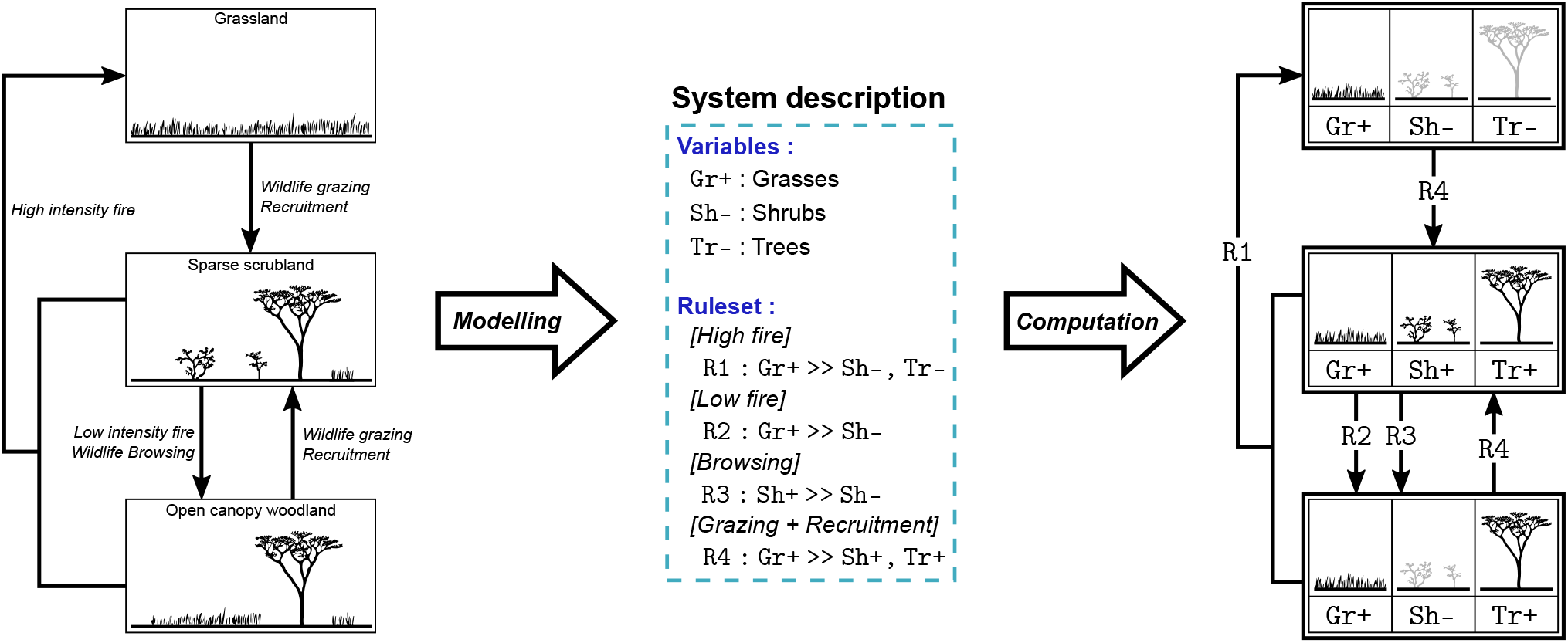
Toy model illustrating the if-then rule modelling. Modelling of the STM of Fig. 1a (left) into a *if-then* rule model (middle: system description) that enables the computation of the STG (right). Each state of the STG is a vector of three Boolean variables described in the “Variables” section of the model description: grasses (`Gr`), shrubs (`Sh`) and trees (`Tr`), noted with + if present and with - if absent. An initial state must be defined to start the computation of the STG, here we chose the grassland after the last high intensity fire [3]. The initial values of the variables are noted next to their symbol in the system description. The “Ruleset” section of the system description encompasses the if-then rules describing the transitions. For example, the first rule `R1` embodies that *if* grasses are present (`Gr+`) *then* (≫) they can fuel a high intensity fire burning down shrubs and trees (`Sh−`, `Tr−`), as grasses resprout first they do not disappear in the fire consequence. This rule corresponds in the STG to the transitions from the middle and bottom states toward the top state. Applying every possible rule to every reachable state computes the STG as depicted on the right, where each transition is labelled with the corresponding rule. A rule is applied if and only if its consequence differs from its condition, thus loops from one state to itself are excluded. Compared to the STM of Fig. 1a, the computed STG is more explicit: there are two transitions between the middle state and the bottom one because two distinct events may lead the system from the former to the latter.

The LBV-model consists of 15 variables, including seven controls (Tab. 1), and 17 rules (S1 Table). Each valuation of the variables describes a state of the Borana ecosystem, that can be classified into vegetation classes [34] (see S2 Table). Each valuation of the controls defines a specific *scenario* for the Borana dynamics (i.e. a combination of altitude and management policies), inspired from historical management and recommendations to limit encroachment [35].

**Table 1.** Variables and controls of the LBV-model. The initial values of the variables are noted next to their name with + if present and − if absent. The initial state of the model describes the grassland vegetation class (only grasses are present) after the last high fire [3]. Each valuation of the controls defines a particular scenario and remains fixed during the dynamic.

| Variable | Description | Control | Description |
| --- | --- | --- | --- |
| Gr+ | Grasses | Alt | Altitude |
| Sh- | Shrubs | Fb | Fire ban |
| Tr- | Trees | Cb | Crop ban |
| Sa- | Tree saplings | Wl | Wildlife presence |
| Cr- | Crops | Ps | Pastoralism |
| Lv- | Livestock | Ig | Intensive grazing |
| Gz- | Wild grazers | BLv | Browsing Livestock |
| Bw- | Wild browsers |  |  |

### 1.2 Model-checking

*Model-checking* is an automated method for the analysis of any dynamical system that can be modelled by states and transitions [40]. Its goal is checking that a given automated system (e.g. hardware or software), modelled as a STG, verifies a given dynamical property, usually written as a temporal logic formula (Fig. 3, in black). In the field of computer science dealing with model-checking, STGs are mainly named *Kripke structures* or *labelled transition systems*, depending if either or both of their states and transitions are labelled. As in systems biology, we will keep calling them state-transition graphs (STGs) for clarity. While model-checking is a wide and active field of research [2], the scope of this paper is limited to exhibiting the potential of model-checking in ecological applications. Thus only one implementation of the model-checking methodology will be illustrated in this paper (Fig. 3, blue italic annotations), while the diversity of the possible implementations will be addressed in the Discussion below.

**Fig 3.**
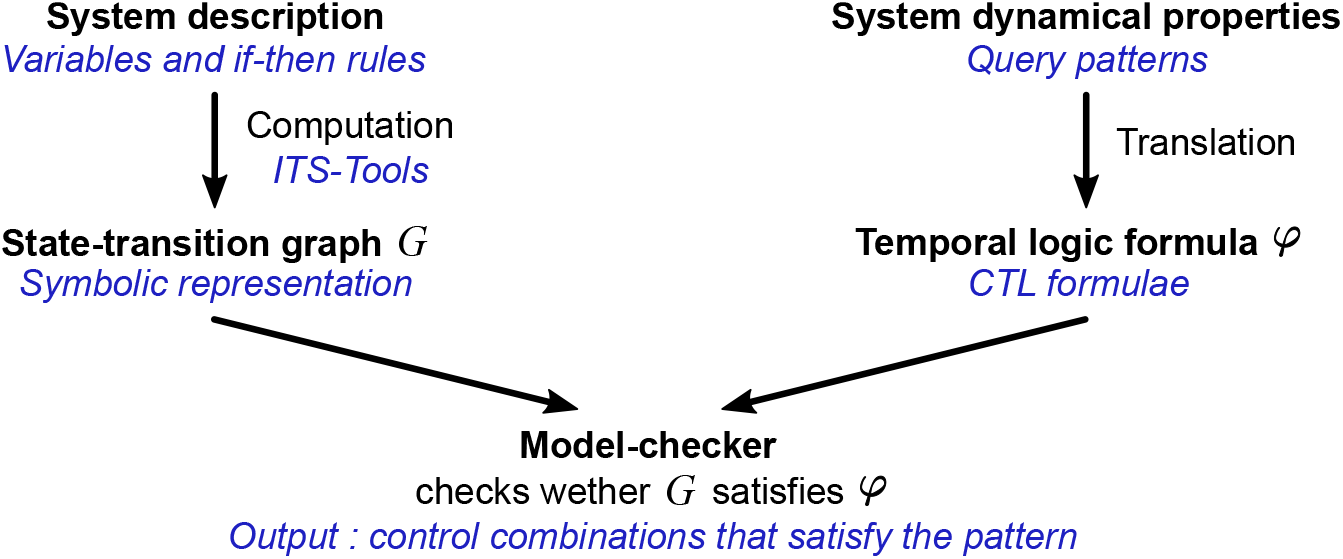
The model-checking methodology. In black the general model-checking outline. In blue italic the implementation described in this article that consists of a particular choice of techniques and tools among the available ones. (Adapted from [40].)

To our knowledge, model-checking has been very scarcely used in ecology, despite its extensive application in systems biology [14]. So far, most formal analyses of STGs in ecology have been limited to graph measures [41] and topology analyses [32, 33, 39]. We identified only a few studies implementing model-checking in ecology [42–45], most modelling an agrosystem as an automated machine in the usual model-checking mindset.

Besides a formal modelling language that enables the precise description of the system of interest (e.g. the if-then rules presented above), model-checking also requires a formal language to express the dynamical properties to be checked, such as temporal logics (Fig. 3).

#### 1.2.1 Expressing properties using Computation Tree Logic (CTL)

*Computation Tree Logic (CTL)* is one of the most popular temporal logic [2] because it is particularly fitted to express properties of branching dynamics with alternative pathways. We chose CTL here because management policies involve choices between alternative pathways, other temporal logics will be presented in Discussion. A CTL formula describes a property over *computation trees*. A computation tree is rooted at a given state of the STG, and its branches are the alternative possible pathways starting from this state (Fig. 4). Let us give two examples of CTL dynamical properties to foster intuition: (1) all the computation tree’s pathways eventually lead to an encroached state, (2) at least one of the computation tree’s pathways maintains grasses. A CTL model-checker discriminates between the states whose computation tree satisfies a given property and those whose computation tree does not. So such a model-checker will check whether the computation tree rooted in each state satisfies a CTL formula or not.

**Fig 4.**
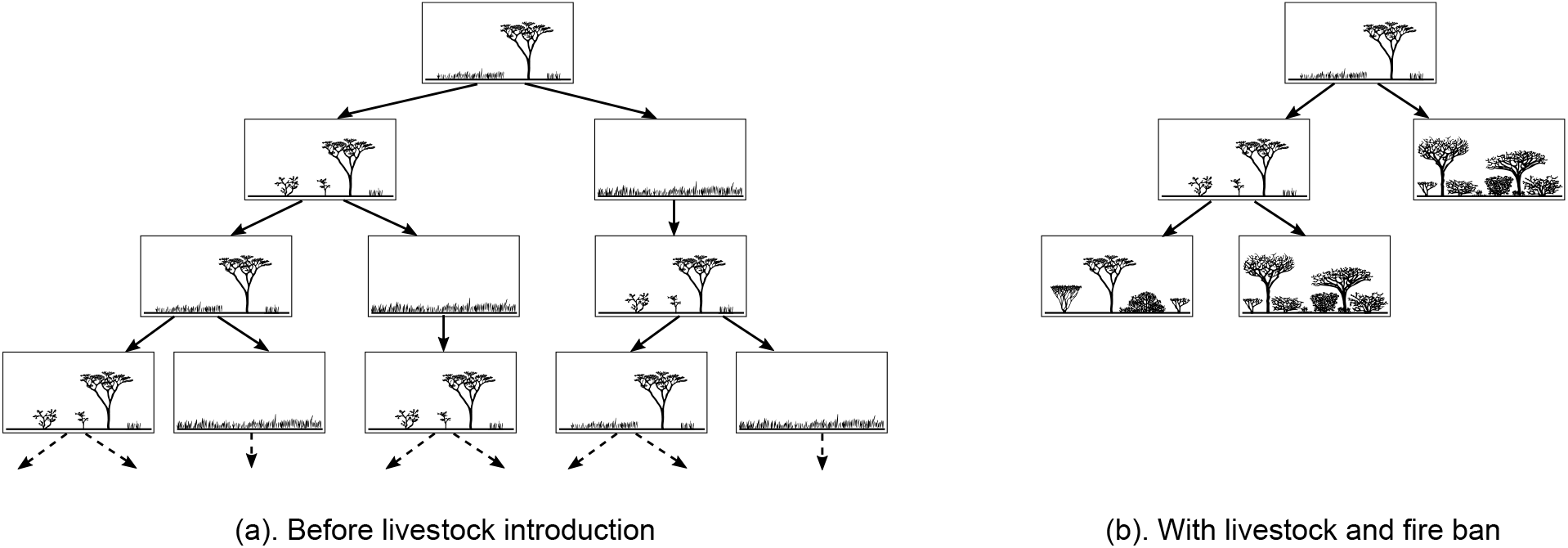
Computation trees rooted in the STMs of Fig. 1a. Each branch descending from the root represents a possible pathway in the STM. (a) The computation tree rooted in the open canopy woodland state of the STM of Fig. 1a. As the pathways are infinite in this STM, the branches of the tree, and thus the tree itself, are also infinite. (b) The computation tree rooted in the open canopy woodland state of the STM of Fig. 1b. The grassland state is not reachable from the open canopy woodland state in this STM, and thus it does not appear in its computation tree. As the pathways are finite in this STM, the branches of the tree are also finite, and thus the tree itself. Formally the branches of a computation tree are usually assumed to be infinite, so that the pathways always continue. In order to tackle this issue, the dead-end leafs of a computation tree can be interpreted as infinite pathways remaining in the same state.

The syntax and semantics of CTL are given in Fig. 5. A *state property p* is a Boolean property mapping over states. For example the presence of shrubs is a state property noted `Sh+`, and in Fig. 2 it is only True (noted by ⊤) over sparse scrubland (middle state). More complex state descriptions are built by combining state properties using the *Boolean logical operators*: not (¬), and (∧), or (∨). For example the encroachment could be defined as the absence of grasses, and the presence of shrubs or trees: `Gr−` ∧ (`Sh+` ∨ `Tr+`). Other Boolean logical operators can be built on top of the three ones above, such as the implication (⟹) that is defined such that *p* ⟹ *q* is equivalent to (¬*p*) ∨ *q*.

**Fig 5.**
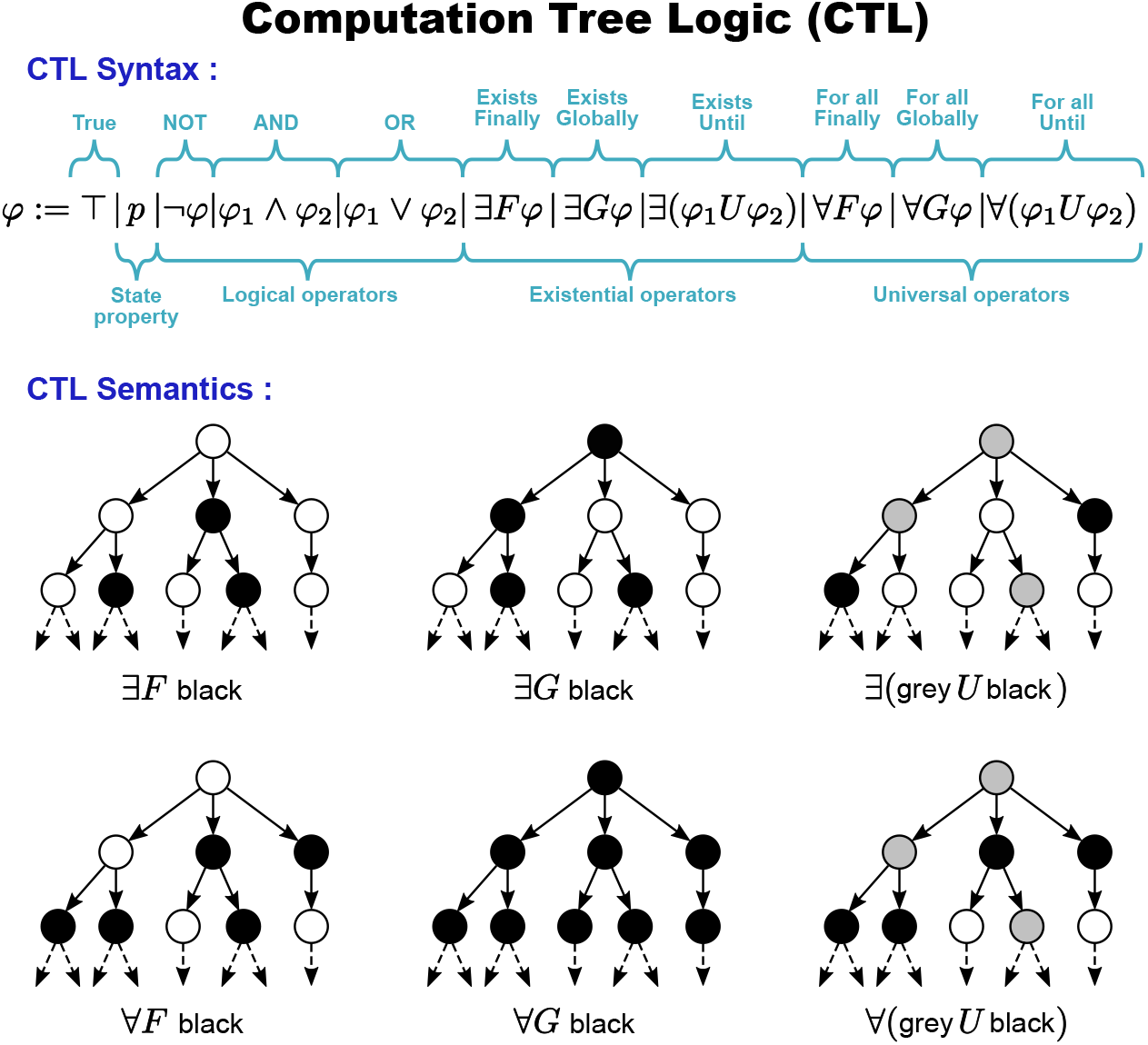
Syntax and semantics of Computation Tree Logic (CTL). The syntax defines how state properties and operators (logical, existential or universal) can be combined into a formula. Besides the well known logical operators, temporal logics introduce temporal operators enabling the expression of temporality (existential and universal operators). The semantics describes the meaning of formulas. The semantics presented here is intuitive and given through example computation trees satisfying basic CTL formulas. (Adapted from [46].) See [2] for a formal semantics of CTL.

The *temporal operators* of CTL are always the combination of two types of operators: first a *quantifier* (∃ or ∀) dealing with non-determinism by quantifying over the pathways starting from a given state, second a *modality* (*F*, *G*, or *U*) specifying the order of events along a pathway. Temporal operators can thus be separated into existential and universal operators. The *existential operators* (∃*F*, ∃*G*, or ∃*U*, see Fig. 5) specify that their modality has to be verified by *at least one branch* of the computation tree (thus by at least one pathway of the STG starting from its root state). The *universal operators* (∀*F*, ∀*G*, or ∀*U*, see Fig. 5) specify that their modality has to be verified by *every branch* of the computation tree (thus by every pathway of the STG starting from its root state). Modality *F* specifies that the property *finally* becomes true at one step of the pathway. Modality *G* specifies that the property is *globally* true all along the pathway. Modality *U* specifies that the left-hand-side property remains true along the pathway *until* the right-hand-side property finally becomes true. Modality *next X* has been omitted from this paper to simplify the presentation.

For example, in computation trees rooted in the Borana STMs (Fig. 4):

- The CTL formula ∃*F* `Tr−` specifies that a state without trees is reachable from the root of the computation tree, which is satisfied in Fig. 4a but not in Fig. 4b.
- The CTL formula ∃*G* `Tr+` specifies that trees are always present along at least one branch of the computation tree, which is satisfied for the left-most branch in Fig. 4a, but not for the other branches.
- The CTL formula ∀*G* `Tr+` specifies that trees are always present all along every branch of the computation tree, which is satisfied in Fig. 4b, but not in Fig. 4a.

Lastly CTL operators can be nested to express even subtler temporal behaviour. For example ∀*G*(∃*F* `Tr−`) specifies that: all along every pathway (∀*G*), the pathway can always branch off to reach a future state (∃*F*) without trees (`Tr−`). While ∀*G*(∃*F* `Tr−`) holds in Fig. 4a, the simpler property ∀*F* `Tr−` does not because trees never disappear in the left-most branch of the computation tree.

Translating an ecological dynamical property (i.e. a description of an ecosystem behaviour) written in English into a CTL formula can turn out to be a delicate exercise for non expert users [47]. One possible way to simplify this task is to provide users with a catalogue mapping patterns of queries to their translations in CTL (Tab. 2) [44, 48].

**Table 2.**
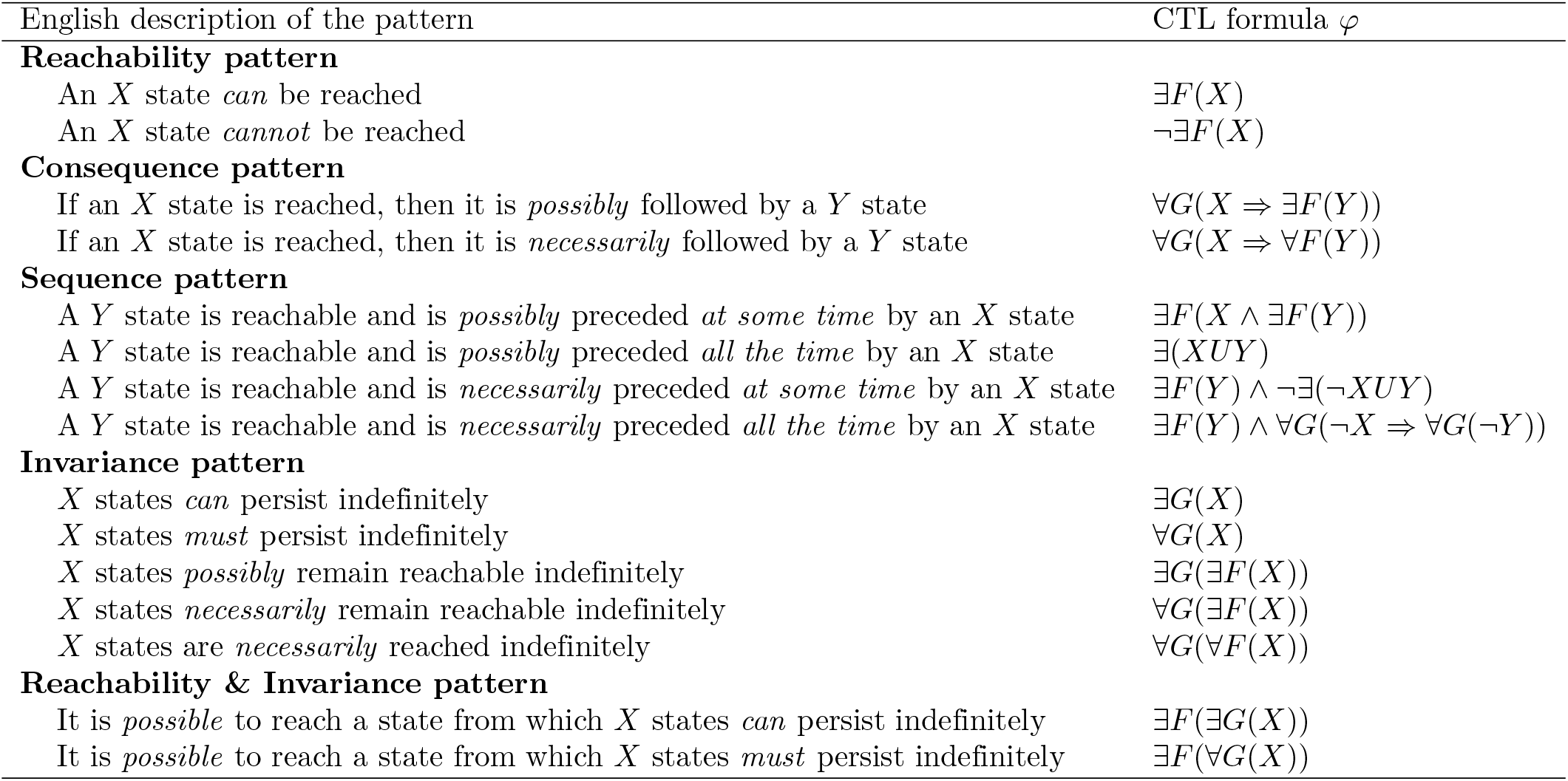
Query patterns of ecological dynamical properties and their translation into CTL formulas. Dynamical properties relevant to ecological systems are gathered into patterns. The patterns are written in English and translated into CTL formulas. *X* and *Y* are place-holders for state properties, e.g. the valuation of a subset of variables. (Adapted from [48].)

| English description of the pattern | CTL formula $\varphi$ |
| --- | --- |
| **Reachability pattern** |  |
| An $X$ state *can* be reached | $\exists F(X)$ |
| An $X$ state *cannot* be reached | $\neg \exists F(X)$ |
| **Consequence pattern** |  |
| If an $X$ state is reached, then it is *possibly* followed by a $Y$ state | $\forall G(X \Rightarrow \exists F(Y))$ |
| If an $X$ state is reached, then it is *necessarily* followed by a $Y$ state | $\forall G(X \Rightarrow \forall F(Y))$ |
| **Sequence pattern** |  |
| A $Y$ state is reachable and is *possibly* preceded *at some time* by an $X$ state | $\exists F(X \wedge \exists F(Y))$ |
| A $Y$ state is reachable and is *possibly* preceded *all the time* by an $X$ state | $\exists(XUY)$ |
| A $Y$ state is reachable and is *necessarily* preceded *at some time* by an $X$ state | $\exists F(Y) \wedge \neg \exists(\neg XUY)$ |
| A $Y$ state is reachable and is *necessarily* preceded *all the time* by an $X$ state | $\exists F(Y) \wedge \forall G(\neg X \Rightarrow \forall G(\neg Y))$ |
| **Invariance pattern** |  |
| $X$ states *can* persist indefinitely | $\exists G(X)$ |
| $X$ states *must* persist indefinitely | $\forall G(X)$ |
| $X$ states *possibly* remain reachable indefinitely | $\exists G(\exists F(X))$ |
| $X$ states *necessarily* remain reachable indefinitely | $\forall G(\exists F(X))$ |
| $X$ states are *necessarily* reached indefinitely | $\forall G(\forall F(X))$ |
| **Reachability & Invariance pattern** |  |
| It is *possible* to reach a state from which $X$ states *can* persist indefinitely | $\exists F(\exists G(X))$ |
| It is *possible* to reach a state from which $X$ states *must* persist indefinitely | $\exists F(\forall G(X))$ |

Note that we have illustrated the semantics of CTL by evaluating formulas with respect to a single root state and its computation tree. More generally, the output of a CTL model-checker is the *set* of all the states of the STG whose computation trees satisfy the formula. This amounts in theory to consider every reachable state of the STG as the root of a computation tree, and to evaluate the formula for each such tree. In practice, model-checkers use much more efficient techniques to obtain their results and avoid this nested enumeration.

We illustrate the model-checking methodology in the Results below. We chose six dynamical queries about the Borana ecosystem covering all five kinds of patterns presented in Tab. 2. Then the scenarios of the LBV-model satisfying each query were computed using the model-checker presented right next.

#### 1.2.2 Our implementation of a scalable model-checker

We instantiated the model-checking methodology (Fig. 3, in black) in a toolkit (Fig. 3, blue italic annotations) including the modelling language based on if-then rules presented in Section 1.1.3, a scalable computation of the STG, and a CTL model-checker (S3 Notebook).

Computation trees do not have to be explicitly constructed in order to check a CTL formula. Even the explicit computation of the STG is not required to solve the CTL model-checking problem. Indeed it can be solved with only a *symbolic* representation of the STG, handling sets of states through an efficient representation that avoids their explicit enumeration [49]. Those symbolic algorithms can mitigate the *combinatorial explosion problem* (i.e. the exponential growth of the number of states with the number of variables) that is inherent to state-based approaches [50].

More precisely, our toolkit transforms the variables and ruleset (Tab. 1 and S1 Table) into a symbolic representation based on *Data Decision Diagram* (*DDD*) [51] provided by ITS-tools [52]. On this basis, we have implemented the CTL symbolic model-checking algorithm from [50]. The resulting model-checker outputs as a DDD the symbolic representation of the states of the STG satisfying a query formula.

In the next Section, we use this toolkit to analyse the LBV-model by determining the scenarios satisfying a given CTL query, i.e the control valuations for which the CTL formula is satisfied in the initial state. Being fully symbolic, our implementation is highly scalable and can handle models with millions of states. This symbolic representation enables the model-checking of a given formula at once for all the possible valuations of the controls of the LBV-model. Thus, the model-checker outputs the DDD holding all the states of all the scenarios satisfying the formula. By extracting only the control variables from this DDD, we are able to observe which combinations of them satisfy a given formula. This information is then transformed into an equivalent Boolean formula that is finally transformed into a *canonical form* using SymPy [53].

## 2 Results

The STG computed from the LBV-model exhibits 1141 states and consists of 2^7^ = 128 disconnected subgraphs, one per control valuation. Thus each subgraph represents the dynamics of the Borana ecosystem under a specific scenario (i.e. a combination of elevation and management policies).

We designed six queries relevant to the Borana ecosystem management and covering all five pattern types (Tab. 2). Those queries are built upon the following state properties:

- Closed canopy woodland is a vegetation class [34] modelled by the presence and absence of some plant variables (see S2 Table for an exhaustive definition of the Borana vegetation classes as state properties):

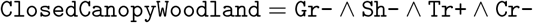
- Encroachment [35] corresponds to the vegetation classes with shrubs or trees but without grasses nor crops (closed canopy woodland, dense scrubland and bushland):

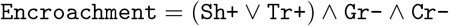
- Subsistence production [54] corresponds to the states with crops or livestock:

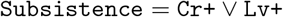

The model-checking outputs the control valuations (scenarios) satisfying each of the queries (Tab. 3). The first query exhibits a straightforward output to the simplest pattern: encroached states are only reachable under pastoralism with intensive grazing. The second query shows that a simple pattern can have a complex output. The remaining queries propose a general survey of the patterns (Tab. 2) with outputs of various complexity. Computing all the model-checking results took about ten seconds on a modern computer (S3 Notebook).

**Table 3.** Model-checker outputs. For each of the six queries we show: (1) its CTL formula and pattern type, (2) its translation into English, (3) the associated output of the model-checker ☑expressed as the scenarios for which the query is satisfied, (4) an English interpretation of this output.

|  |  |
| --- | --- |
| 1) | **Reachability pattern:** $\exists F \text{Encroachment}$<br>An encroached state can be reached.<br>$\checkmark \text{ Ps+} \wedge \text{Ig+}$<br>*Encroachment can only be reached under the scenarios encompassing pastoralism Ps+ with intensive grazing Ig+.* |
| 2) | **Reachability pattern:** $\exists F \text{ClosedCanopyWoodland}$<br>Closed Canopy Woodland can be reached.<br>$\checkmark \text{ Ps+} \wedge \text{Ig+} \wedge (\text{Alt+} \vee \text{Fb-} \vee \text{Wl+} \vee \text{BLv+})$<br>*Closed Canopy Woodland can only be reached under pastoralism Ps+ with intensive grazing Ig+ and with at least one of the following factors: high altitude Alt+, no fire ban Fb-, presence of wildlife Wl+, browsing livestock BLv+.* |
| 3) | **Reachability + Consequence pattern:** $(\exists F \text{Encroachment}) \wedge (\forall G(\text{Encroachment} \Rightarrow \exists F \neg \text{Encroachment}))$<br>An encroached state is reachable, and whenever it is reached it is possibly followed by an unencroached state.<br>$\checkmark \text{ Ps+} \wedge \text{Ig+} \wedge \text{Alt+} \wedge \text{Cb-}$<br>*If an encroached state is reachable (see output Ps+ $\wedge$ Ig+ from query 1) and if the system is at high altitude Alt+ with crops allowed Cb-, then whenever an encroached state is reached it is possibly followed by an unencroached state, i.e. encroachment is reversible.* |
| 4) | **Sequence pattern:** $(\exists F \text{Encroachment}) \wedge \neg \exists (\neg (\text{Lv+} \vee \text{Gz+}) U \text{Encroachment})$<br>An encroached state is reachable and is necessarily preceded at some time by a state with grazers.<br>$\checkmark \text{ Ps+} \wedge \text{Ig+}$<br>*If an encroached state is reachable (see output Ps+ $\wedge$ Ig+ from query 1) then it is necessarily preceded by the presence of livestock or the presence of wild grazers.* |
| 5) | **Invariance pattern:** $\forall G(\exists F \text{Subsistence})$<br>Subsistence states necessarily remain reachable indefinitely.<br>$\checkmark (\text{Ps+} \wedge \text{Ig-}) \vee (\text{Alt+} \wedge \text{Cb-} \wedge \text{Ps+}) \vee (\text{Alt+} \wedge \text{Cb-} \wedge \text{Wl+})$<br>*There are three sets of scenarios where subsistence remains reachable whatever happens : (1) under pastoralism Ps+ without intensive grazing Ig-, (2) at high altitude Alt+ with crops allowed Cb- and with pastoralism Ps+, or (3) at high altitude Alt+ with crops allowed Cb- and with wildlife Wl+.* |
| 6) | **Reachability & Invariance pattern:** $\exists F(\exists G \text{Subsistence})$<br>It is possible to reach a state from which the subsistence can persist indefinitely.<br>$\checkmark (\text{Ps+} \wedge \text{BLv+}) \vee (\text{Alt-} \wedge \text{Ps+}) \vee (\text{Fb+} \wedge \text{Cb+} \wedge \text{Ps+} \wedge \text{Ig-})$<br>*There are three sets of scenarios where it is possible to reach a state from which subsistence can persist indefinitely: (1) under pastoralism Ps+ with browsing livestock BLv+, (2) at low altitude Alt- with pastoralism Ps+, or (3) with fire banned Fb+ as well as crops Cb+ and with pastoralism Ps+ but without intensive grazing Ig-.* |

In order to compare under various scenarios the LBV-model with Liao’s STMs [(Liao et al. 2018b, Fig. 5)](https://doi.org/10.1016/j.agee.2017.10.009), we split the states of the LBV-model between the vegetation classes. Thus the STGs drawn below aggregate the states of the LBV-model into vegetation classes. An edge is drawn between two vegetation classes if and only if there is at least one transition in the STG of the LBV-model from a state of the first vegetation class to a state of the second vegetation class. Those edges are labelled with the corresponding transitions of the LBV-model’s STG. The initial states of the LBV-model are chosen accordingly with the corresponding scenarios.

Nodes (vegetation classes) decorated with a downward triangle contain initial states, nodes decorated with a circle contain cyclic pathways in the LBV-model STG. The nodes colors are chosen arbitrarily.

The queries outputs (Tab. 3) exhibit how the model-checking methodology could help better understanding the Borana vegetation pathways and choosing adequate management policies. The first two queries describe the scenarios enabling encroachment. The third query describes the scenarios making encroachment reversible. The fourth query shows that grazing is a necessary condition for encroachment. The fifth query describes the scenarios enabling at least chronic subsistence (food is not constantly but only regularly reachable). The sixth query describes the scenarios enabling persistent subsistence.

The experiments on the LBV-model should be read as a proof-of-concept illustrating the model-checking methodology we promote, rather than as a truthful modelling of an ecological system. Indeed, validating the LBV-model is out of the scope of this paper, and the LBV-model could certainly be improved to give a more accurate representation of the Borana vegetation dynamics. However, the LBV-model is a faithful representation of what an ecological STG looks like, and has been developed to exhibit what model-checking can provide to ecology. Thus the model-checking results should be read as a demonstration of the insights offered by model-checking, rather than as actual recommendations for the Borana vegetation management.

## 3 Discussion

### 3.1 Model-checking ecological state-transition graphs

Model-checking performs efficient and automated explorations of the temporal behaviour of ecological STGs, answering a recently expressed interest for such tools [9, 10]. Mathematically defining the dynamical properties of the system with the help of temporal logic can help removing ambiguity in definitions and thus improve reproducibility. Moreover since model-checking is automated, it can process STGs that are too large to be examined by hand, thus providing more precise and complex descriptions of ecosystem dynamics. Indeed, CTL model-checking has been applied in systems biology to STG models made of hundreds of variables [55].

Even on our LBV-model that yields “only” 1141 states, answering questions like “*under what scenarios subsistence necessarily remains regularly reachable*” (Tab. 3, query 5) would probably be hardly feasible without resorting to model-checking. Not only this question actually corresponds to a not-so-simple CTL property, but its answer is also surprisingly complex. A human examination may certainly detect the importance of pastoralism, crop ban and altitude, but would most likely fail to accurately relate them within a reasonable amount of time. On the other hand, model-checking is fully automated and provides the exact answer in a matter of seconds. Human work is then limited to the conception of temporal logic formulas, which is beneficial to science reproducibility.

To illustrate the model-checking of ecological STGs, we have instantiated its general outline (Fig. 3, in black) with a particular choice of methods and tools (Fig. 3, blue italic annotations). This particular instance is only illustrative and will be discussed in the following. Note that a computational model, i.e. the system description and the computation step (Fig. 3), is not mandatory in the model-checking methodology. Indeed complex STGs can be found directly in empirical studies [5, 56], without being computed from an underlying mathematical system description. Hence model-checking is not only a tool for the analysis of theoretical models, but could also assist the automated investigation of empirical data. Bearing this in mind, we will now focus on the various potential implementations of the system description (Fig. 3, left-most half).

### 3.2 Choosing a system description

The LBV-model is built from if-then rules (Fig. 2), a methodology previously proposed to model ecosystem dynamics from expert knowledge [31, 37–39]. STGs can also be built from differential equations [9] or Boolean networks [32, 57], with possible bridges between both [33]. Model-checking tools manipulating biological networks have been designed in the field of systems biology [58, 59], for example GINsim [60] handling Boolean networks.

We have chosen here a description of the system based on events (if-then rules) because it looks suited to the available data in the STM literature [7], i.e. the list of the transitions between states and their main drivers. Descriptions based on Boolean networks are centred on interaction networks, following for example the methodology built in systems biology to study regulatory networks [61, 62]. Another difference between both modelling approaches is that event-based modelling can express non-determinism at the level of rules. Indeed, we may write rules with the same conditions but leading to contradictory consequences, for instance to represent partial knowledge about the conditions that drive a system. However, this is not possible with Boolean networks where each variable is updated in a deterministic way through a fixed function. In such a setting, non-determinism may only arise at the level of the STG from the order in which variables are updated, including the possibility or not to update them simultaneously. In particular, the non-determinism of the STG of Fig. 2 could not be obtained from a Boolean network, hence our choice of a if-then rule model.

Lastly the LBV-model is built upon Boolean variables, which are either present or absent. Yet in general, the model-checking methodology can be applied to any discrete-state model description computing into a STG. Thus variables can be multivalued, which is typically used in systems biology to model phenomena where a reactant regulates distinct reactions that occur at distinct thresholds. In the Borana ecosystem, fire is rare when woody plant cover is above a threshold of 40% [35, 63], thus trees would better be described as multivalued rather than as Boolean:

Tr ∈ {*none, low, high*} corresponding respectively to 0%, < 40% and ≥40%.

Computer science provides with a large range of modelling formalisms that cover these needs and compute STGs. Some are used in systems biology and thus may be suitable for ecology as well [15, 59].

### 3.3 Choosing a temporal logic

As for the system description, numerous possibilities exist to translate the dynamical properties of a system into formulas in one of the existing temporal logics (Fig. 3, right-most half). In this paper we have introduced a chosen set of patterns and their translation (Tab. 2) into one very popular temporal logic: the Computation Tree Logic (CTL) that expresses branching properties between alternative pathways (Fig. 5). This is suitable to represent the properties of the Borana ecosystem dynamics because management actions can be represented as choices between alternative pathways.

Another very popular temporal logic is Linear-time Temporal Logic (LTL) [2] that expresses complex properties about a single pathway (hence its linear representation of time) by nesting temporal modalities. For example LTL could be used to validate models because the available observations often consist of particular pathways [64]. CTL and LTL together are included into the richer temporal logic CTL* [2] that handles both branchings and complex pathway properties. However the model-checking of CTL* is computationally much harder than the model-checking of either CTL or LTL, and thus it is less used in practice.

Instead of defining management policies as controls (i.e. invariant variables) like in the LBV-model (Tab. 1), they can label the transitions they enable. As management policies are no longer control variables, the number of states is reduced, as well as the computational complexity. Moreover management policies can then vary along pathways, enabling the assessment of adaptive management strategies. Action Restricted CTL (ARCTL) [65] is an extension of CTL that deals with such transition labels and can thus handle the branchings induced by changes in management policies. ARCTL has been applied successfully in systems biology [17].

Finally, choosing the next transition can be represented as a game between agents of the ecosystem (in the game theory framework). In this perspective, each transition results from a coalition of some agents that “collaborate” to perform a change in the system. Alterning-time Temporal Logic (ATL) [66] is an extension of CTL that handles queries about the outcomes of such games. This representation could help collaborative decision making which is at the heart of the STM framework [7].

### 3.4 Choosing a modelling and analysis framework

More features may be wanted from the modelling and properties languages. For example, when the durations of the transitions between states are available and of interest [67], they may be incorporated into the model description resulting into timed automata [44, 68, 69] or time Petri net [70]. Several temporal logics have been designed to handle quantitative durations of transitions [71], for example Timed CTL (TCTL) [72] an extension of CTL (see [45] for an application on agrosystems), or Metric Temporal Logic (MTL) [73] an extension of LTL. Similarly when the probabilities of the transitions are available, they may be incorporated into the model description resulting into discrete time Markov chain, for example [74], and the model-checking methodology can be adapted to handle stochastic phenomena [75].

Such extensions lead to hybrid systems that mix discrete states with continuous information such as time or probabilities. On the one hand, they are appealing because they offer a wider range of modelling features than simpler purely discrete systems. But on the other hand, they turn out to be computationally much harder. Thus they offer fewer analysis capabilities and fewer software tools exist to support them. In general, the more expressive a particular setting is, the less it can be analysed (because of computational complexity and tools availability), and some questions even become undecidable (i.e. no algorithm can answer them).

So, the choice of a particular system description and temporal logic results from a trade-off between the features one needs (or would like) to include and the analysis one needs (or would like) to perform. Features that cannot be represented will be abstracted away, while analyses that cannot be performed must be given up. Thus the modelling and analysis framework must be chosen accordingly to the available data and to the targeted question.

For instance, the if-then rules presented in this paper form quite a general Boolean setting for the systems description, while opening a wide range of analysis possibilities. In particular, model-checking using any of the temporal logics presented in the previous section can be envisaged. In this case, the compromise is in favour of the analysis power while systems have to be represented in abstract ways. The success of Boolean networks in systems biology probably shows that there is already much to learn from abstract qualitative models [15].

## 4 Conclusion

This article promotes the model-checking of ecological state-transition graphs. Although STGs are common in ecology, from state-and-transition models to assembly graphs, the model-checking methodology remains widely unused. Yet model-checking proved in systems biology to be a valuable automated tool for the analysis of STGs, resulting into many already available software packages. Model-checking can be performed on both theoretical models and empirical data, as long as they sum up into a STG. The model-checking methodology encompasses a broad range of concepts and tools, thus its implementation can be fitted to the specific features of the system under study (for example durations or probabilities labelling the transitions). In addition to the automated analysis of ecological STGs, we believe that definitions based on temporal logics would help clarifying and comparing the various concepts used in the related fields of ecology.

## Supporting information

**S1 Table.** Ruleset of the LBV-model. The 17 if-then rules describing the vegetation dynamics in Borana. The left part of a rule is its condition (if …), the right part is its consequence (then …). For example the first rule `R1` states that *if* fire is allowed (`Fb−`) and grasses are present (`Gr+`), *then* (≫) a low intensity fire can occur, resulting in the burning of shrubs and saplings (`Sh−`, `Sa−`) and in the animals escape (`Lv−`, `Gz−`, `Bw−`), as grasses resprout first they do not disappear in the fire consequence. This ruleset was inspired from the STM literature on Borana [3, 34–36].

|  | Description | Condition (if ...) | Consequence (then ...) |
| --- | --- | --- | --- |
| R1 | *Low fire* | Fb-, Gr+ >> | Sh-, Sa-, Lv-, Gz-, Bw- |
| R2 | *High fire* | Fb-, Gr+ >> | Sh-, Tr-, Sa-, Lv-, Gz-, Bw- |
| R3 | *Trees* | Sa+ >> | Tr+ |
| R4 | *Grass* | Sh-, Tr-, Sa-, Cr- >> | Gr+ |
| R5 | *CCW* | Alt+, Fb+, Gr-, Sa+ >> | Sh-, Tr+ |
| R6 | *Bushland* | Alt-, Sh+, Tr- >> | Sa- |
| R7 | *Grazing* | Gr+, Lv+ >> | Sh+, Sa+ |
| R8 | *Grazing* | Gr+, Gz+ >> | Sh+, Sa+ |
| R9 | *Intensive grazing* | Ig+, Gr+, Lv+ >> | Gr-, Lv-, Gz- |
| R10 | *Grazers* | Wl+, Gr+, Lv- >> | Gz+ |
| R11 | *Livestock* | Ps+, Gr+ >> | Lv+, Gz-, Bw- |
| R12 | *Browsing* | Bw+ >> | Gr+, Sh-, Sa-, Bw- |
| R13 | *Cattle Browsing* | BLv+, Lv+ >> | Gr+, Sh-, Sa-, Bw- |
| R14 | *Browsers* | Wl+, Sh+, Lv- >> | Bw+ |
| R15 | *Browsers* | Wl+, Sa+, Lv- >> | Bw+ |
| R16 | *Crops* | Alt+, Cb-, Tr+ >> | Gr-, Sh-, Sa-, Cr+, Lv-, Gz-, Bw- |
| R17 | *Crops* | Cr+ >> | Gr+, Cr- |

**S2 Table.**
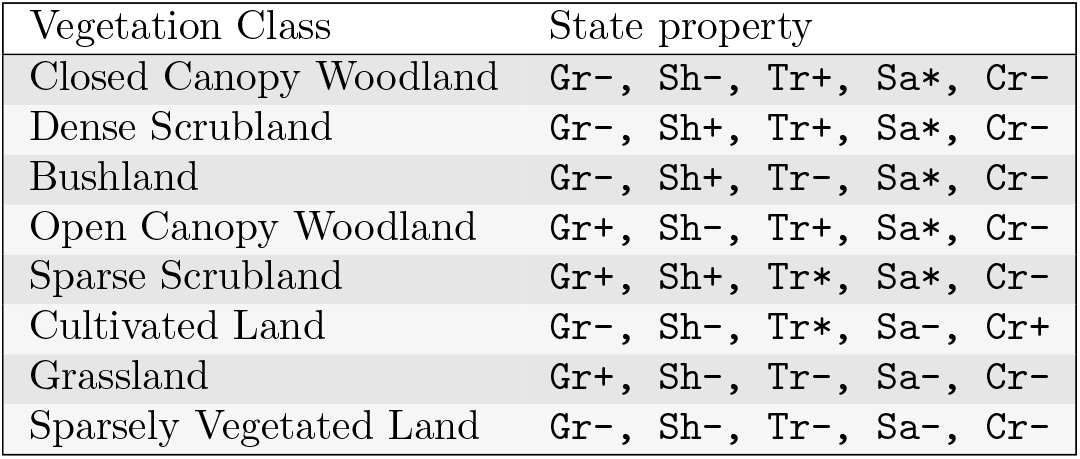
Borana vegetation classes as state properties. Borana vegetation classes [34] translated into state properties (presence or absence of vegetation variables). A variable always present in the vegetation class is noted with “+”, a variable always absent is noted with “`−`”, while a variable whose presence is variable in the vegetation class is noted with “`*`”. For example the closed canopy woodland is described by the state property stating that trees are present, grasses, shrubs and crops are absent, while saplings can be either present or absent. Note that grasses are considered functionally present in the sparse scrubland class although covering only between 10% and 30% of the surface [34], indeed both fire and grazing occur in sparse scrublands [35].

**S3 Notebook. Python notebook encompassing the LBV-model analysis.** Zip archive containing: (1) “`README`” is a text file explaining how to install our implementation of the model-checking methodology (Sec. 1.2.2), (2) “`LBV model.rr`” is a text file containing the system description of the LBV-model (Tab. 1 and S1 Table), “`S3 notebook.ipynb`” is a Jupyter notebook covering the LBV-model analysis (Sec. 2) and a comparison between the LBV-model and Liao’s STMs (Fig. 1), “`S3 notebook.html`” is a static HTML preview of this notebook.

## Acknowledgements

We warmly thank Yann Thierry-Mieg for the fruitful discussions about the ITS-tools framework, as well as Chuan Liao for his kind permission to redraw his STM.

## References

1. Clements FE. Plant succession: an analysis of the development of vegetation. 242. Carnegie Institution of Washington; 1916.

2. Clarke EM, Jr, Grumberg O, Kroening D, Peled D, Veith H. Model Checking. 2nd ed. Cyber Physical Systems Series. Cambridge, MA, USA: MIT Press; 2018.

3. Liao C, Clark PE. Rangeland vegetation diversity and transition pathways under indigenous pastoralist management regimes in southern Ethiopia. Agriculture, Ecosystems & Environment. 2018;252:105–113. doi:10.1016/j.agee.2017.10.009.

4. Chen HY, Popadiouk RV. Dynamics of North American boreal mixedwoods. Environmental Reviews. 2002;10(3):137–166. doi:10.1139/a02-007.

5. Warren PH, Law R, Weatherby AJ. Mapping the Assembly of Protist Communities in Microcosms. Ecology. 2003;84(4):1001–1011. doi:10.1890/0012-9658(2003)084[1001:MTAOPC]2.0.CO;2.

6. Westoby M, Walker B, Noy-Meir I. Opportunistic management for rangelands not at equilibrium. Journal of Range Management. 1989;42(4):266–274. doi:10.2307/3899492.

7. Bestelmeyer BT, Ash A, Brown JR, Densambuu B, Fernández-Giménez M, Johanson J, et al. State and Transition Models: Theory, Applications, and Challenges. In: Briske DD, editor. Rangeland Systems: Processes, Management and Challenges. Springer Series on Environmental Management. Cham: Springer International Publishing; 2017. p. 303–345. Available from: https://doi.org/10.1007/978-3-319-46709-2_9.

8. Hang-Kwang L, Pimm SL. The Assembly of Ecological Communities: A Minimalist Approach. Journal of Animal Ecology. 1993;62(4):749–765. doi:10.2307/5394.

9. Servań CA, Allesina S. Tractable models of ecological assembly. Ecology Letters. 2021;24(5):1029–1037. doi:10.1111/ele.13702.

10. Walker B, Westoby M. Past, present and future of state and transition language. Rangeland Journal. 2020;42(1):71–72. doi:10.1071/RJ20020.

11. Schröter C, Schwoon S, Esparza J. The Model-Checking Kit. In: van der Aalst WMP, Best E, editors. Applications and Theory of Petri Nets 2003. Lecture Notes in Computer Science. Berlin, Heidelberg: Springer; 2003. p. 463–472.

12. Kordon F, Bouvier P, Garavel H, Hillah LM, Hulin-Hubard F, Amat N, et al. Complete Results for the 2020 Edition of the Model Checking Contest; 2021. Available from: http://mcc.lip6.fr/2021/results.php.

13. Wang RS, Saadatpour A, Albert R. Boolean modeling in systems biology: an overview of methodology and applications. Physical Biology. 2012;9(5):055001. doi:10.1088/1478-3975/9/5/055001.

14. Brim L, Češka M, Šafránek D. Model Checking of Biological Systems. In: Bernardo M, de Vink E, Di Pierro A, Wiklicky H, editors. Formal Methods for Dynamical Systems. SFM 2013. Lecture Notes in Computer Science. Berlin, Heidelberg: Springer; 2013. p. 63–112. Available from: https://doi.org/10.1007/978-3-642-38874-3_3.

15. Bartocci E, Lió P. Computational Modeling, Formal Analysis, and Tools for Systems Biology. PLOS Computational Biology. 2016;12(1):e1004591. doi:10.1371/journal.pcbi.1004591.

16. Batt G, Ropers D, de Jong H, Geiselmann J, Mateescu R, Page M, et al. Validation of qualitative models of genetic regulatory networks by model checking: analysis of the nutritional stress response in Escherichia coli. Bioinformatics. 2005;21(suppl 1):i19–i28. doi:10.1093/bioinformatics/bti1048.

17. Abou-Jaoudé W, Monteiro PT, Naldi A, Grandclaudon M, Soumelis V, Chaouiya C, et al. Model Checking to Assess T-Helper Cell Plasticity. Frontiers in Bioengineering and Biotechnology. 2015;2:86. doi:10.3389/fbioe.2014.00086.

18. Traynard P, Fauré A, Fages F, Thieffry D. Logical model specification aided by model-checking techniques: application to the mammalian cell cycle regulation. Bioinformatics. 2016;32(17):i772–i780. doi:10.1093/bioinformatics/btw457.

19. Béal J, Pantolini L, Noël V, Barillot E, Calzone L. Personalized logical models to investigate cancer response to BRAF treatments in melanomas and colorectal cancers. PLOS Computational Biology. 2021;17(1):e1007900. doi:10.1371/journal.pcbi.1007900.

20. Young TP, Chase JM, Huddleston RT. Community succession and assembly comparing, contrasting and combining paradigms in the context of ecological restoration. Ecological Restoration. 2001;19(1):5–18. doi:10.3368/er.19.1.5.

21. Chang C, HilleRisLambers J. Integrating succession and community assembly perspectives. F1000Research. 2016;5:F1000 Faculty Rev–2294. doi:10.12688/f1000research.8973.1.

22. May RM. Network structure and the biology of populations. Trends in Ecology & Evolution. 2006;21(7):394–399. doi:10.1016/j.tree.2006.03.013.

23. Pilosof S, Porter MA, Pascual M, Kéfi S. The multilayer nature of ecological networks. Nature Ecology & Evolution. 2017;1(4):1–9. doi:10.1038/s41559-017-0101.

24. Patten BC. A primer for ecological modeling and simulation with analog and digital computers. In: Systems Analysis and Simulation in Ecology (ed. BC Patten) vol. I. vol. 1. Academic Press; 1971. p. 3–121.

25. Londo G. Successive mapping of dune slack vegetation. Vegetatio. 1974;29(1):51–61. doi:10.1007/BF02390895.

26. Bergeron Y, Chen HYH, Kenkel NC, Leduc AL, Macdonald SE. Boreal mixedwood stand dynamics: ecological processes underlying multiple pathways. Forestry Chronicle. 2014;90(2):202–213.

27. Caudle D. Interagency ecological site handbook for rangelands. US Department of the Interior, Bureau of Land Management; 2013. Available from: https://jornada.nmsu.edu/sites/jornada.nmsu.edu/files/InteragencyEcolSiteHandbook.pdf.

28. Phillips JD, Van Dyke C. State-and-transition models in geomorphology. CATENA. 2017;153:168–181. doi:10.1016/j.catena.2017.02.009.

29. Tittonell P. Assessing resilience and adaptability in agroecological transitions. Agricultural Systems. 2020;184:102862. doi:10.1016/j.agsy.2020.102862.

30. Salles P, Bredeweg B. Modelling population and community dynamics with qualitative reasoning. Ecological Modelling. 2006;195(1):114–128. doi:10.1016/j.ecolmodel.2005.11.014.

31. Mao Z, Centanni J, Pommereau F, Stokes A, Gaucherel C. Maintaining biodiversity promotes the multifunctionality of social-ecological systems: holistic modelling of a mountain system. Ecosystem Services. 2021;47:101220. doi:10.1016/j.ecoser.2020.101220.

32. Campbell C, Yang S, Albert R, Shea K. A network model for plant–pollinator community assembly. Proceedings of the National Academy of Sciences. 2011;108(1):197–202. doi:10.1073/pnas.1008204108.

33. Robeva R, Murrugarra D. The spruce budworm and forest: a qualitative comparison of ODE and Boolean models. Letters in Biomathematics. 2016;3(1):75–92. doi:https://doi.org/10.1080/23737867.2016.1197804.

34. Liao C, Clark PE, DeGloria SD. Bush encroachment dynamics and rangeland management implications in southern Ethiopia. Ecology and Evolution. 2018;8(23):11694–11703. doi:10.1002/ece3.4621.

35. Liao C, Agrawal A, Clark PE, Levin SA, Rubenstein DI. Landscape sustainability science in the drylands: mobility, rangelands and livelihoods. Landscape Ecology. 2020;35(11):2433–2447. doi:10.1007/s10980-020-01068-8.

36. Liao C. Complexity In The Open Grazing System: Rangeland Ecology, Pastoral Mobility And Ethnobotanical Knowledge In Borana, Ethiopia [PhD Thesis]. Cornell University; 2016. Available from: https://hdl.handle.net/1813/43578.

37. Rykiel EJ. Artificial intelligence and expert systems in ecology and natural resource management. Ecological Modelling. 1989;46(1):3–8. doi:10.1016/0304-3800(89)90066-5.

38. Starfield AM. Qualitative, Rule-Based Modeling. BioScience. 1990;40(8):601–604. doi:10.2307/1311300.

39. Gaucherel C, Pommereau F. Using discrete systems to exhaustively characterize the dynamics of an integrated ecosystem. Methods in Ecology and Evolution. 2019;10(9):1615–1627. doi:10.1111/2041-210X.13242.

40. Clarke EM, Henzinger TA, Veith H. Introduction to Model Checking. In: Handbook of Model Checking. Cham: Springer International Publishing; 2018. p. 1–26. Available from: https://doi.org/10.1007/978-3-319-10575-8_1.

41. Phillips JD. The structure of ecological state transitions: Amplification, synchronization, and constraints in responses to environmental change. Ecological Complexity. 2011;8(4):336–346. doi:10.1016/j.ecocom.2011.07.004.

42. Scopélitis J, Andréfouët S, Largouët C. Modelling coral reef habitat trajectories: Evaluation of an integrated timed automata and remote sensing approach. Ecological Modelling. 2007;205(1):59–80. doi:10.1016/j.ecolmodel.2007.02.011.

43. Hélias A, Guerrin F, Steyer JP. Using timed automata and model-checking to simulate material flow in agricultural production systems—Application to animal waste management. Computers and Electronics in Agriculture. 2008;63(2):183–192. doi:10.1016/j.compag.2008.02.008.

44. Largouët C, Cordier MO, Bozec YM, Zhao Y, Fontenelle G. Use of timed automata and model-checking to explore scenarios on ecosystem models. Environmental Modelling & Software. 2012;30:123–138. doi:10.1016/j.envsoft.2011.08.005.

45. Cordier MO, Largouët C, Zhao Y. Model-Checking an Ecosystem Model for Decision-Aid. In: 2014 IEEE 26th International Conference on Tools with Artificial Intelligence; 2014. p. 539–543.

46. Baier C, Katoen JP. Principles of Model Checking. Cambridge, MA, USA: MIT Press; 2008.

47. Dwyer MB, Avrunin GS, Corbett JC. Patterns in property specifications for finite-state verification. In: Proceedings of the 21st international conference on Software engineering. ICSE ‘99. New York, NY, USA: Association for Computing Machinery; 1999. p. 411–420. Available from: https://doi.org/10.1145/302405.302672.

48. Monteiro PT, Ropers D, Mateescu R, Freitas AT, de Jong H. Temporal logic patterns for querying dynamic models of cellular interaction networks. Bioinformatics. 2008;24(16):i227–i233. doi:10.1093/bioinformatics/btn275.

49. Bryant RE. Binary Decision Diagrams. In: Clarke EM, Henzinger TA, Veith H, Bloem R, editors. Handbook of Model Checking. Cham: Springer International Publishing; 2018. p. 191–217. Available from: https://doi.org/10.1007/978-3-319-10575-8_7.

50. Burch JR, Clarke EM, McMillan KL, Dill DL, Hwang LJ. Symbolic model checking: 1020 States and beyond. Information and Computation. 1992;98(2):142–170. doi:10.1016/0890-5401(92)90017-A.

51. Couvreur JM, Encrenaz E, Paviot-Adet E, Poitrenaud D, Wacrenier PA. Data Decision Diagrams for Petri Net Analysis. In: Esparza J, Lakos C, editors. Application and Theory of Petri Nets 2002. Lecture Notes in Computer Science. Berlin, Heidelberg: Springer; 2002. p. 101–120.

52. Thierry-Mieg Y. Symbolic Model-Checking Using ITS-Tools. In: Baier C, Tinelli C, editors. Tools and Algorithms for the Construction and Analysis of Systems. Lecture Notes in Computer Science. Berlin, Heidelberg: Springer; 2015. p. 231–237.

53. Meurer A, Smith CP, Paprocki M, Čertík O, Kirpichev SB, Rocklin M, et al. SymPy: symbolic computing in Python. PeerJ Computer Science. 2017;3:e103. doi:10.7717/peerj-cs.103.

54. Wharton CR. Subsistence Agriculture: Concepts and Scope. In: Subsistence Agriculture & Economic Development. Routledge; 1969. p. 12–20. Available from: https://doi.org/10.4324/9781315130408.

55. Chabrier N, Fages F. Symbolic Model Checking of Biochemical Networks. In: Priami C, editor. Computational Methods in Systems Biology. Lecture Notes in Computer Science. Berlin, Heidelberg: Springer; 2003. p. 149–162.

56. Barrio IC, Hik DS, Thórsson J, Svavarsdóttir K, Marteinsdóttir B, Jońsdóttir IS. The sheep in wolf’s clothing? Recognizing threats for land degradation in Iceland using state-and-transition models. Land Degradation & Development. 2018;29(6):1714–1725. doi:10.1002/ldr.2978.

57. LaBar T, Campbell C, Yang S, Albert R, Shea K. Global versus local extinction in a network model of plant–pollinator communities. Theoretical Ecology. 2013;6(4):495–503. doi:10.1007/s12080-013-0182-8.

58. Carrillo M, Gońgora PA, Rosenblueth D. An overview of existing modeling tools making use of model checking in the analysis of biochemical networks. Frontiers in Plant Science. 2012;3:155. doi:10.3389/fpls.2012.00155.

59. Naldi A, Monteiro PT, Müssel C, the Consortium for Logical Models and Tools, Kestler HA, Thieffry D, et al. Cooperative development of logical modelling standards and tools with CoLoMoTo. Bioinformatics. 2015;31(7):1154–1159. doi:10.1093/bioinformatics/btv013.

60. Chaouiya C, Naldi A, Thieffry D. Logical Modelling of Gene Regulatory Networks with GINsim. In: van Helden J, Toussaint A, Thieffry D, editors. Bacterial Molecular Networks: Methods and Protocols. Methods in Molecular Biology. New York, NY: Springer; 2012. p. 463–479. Available from: https://doi.org/10.1007/978-1-61779-361-5_23.

61. Abou-Jaoudé W, Traynard P, Monteiro PT, Saez-Rodriguez J, Helikar T, Thieffry D, et al. Logical Modeling and Dynamical Analysis of Cellular Networks. Frontiers in Genetics. 2016;7. doi:10.3389/fgene.2016.00094.

62. Schwab JD, Kühlwein SD, Ikonomi N, Kühl M, Kestler HA. Concepts in Boolean network modeling: What do they all mean? Computational and Structural Biotechnology Journal. 2020;18:571–582. doi:10.1016/j.csbj.2020.03.001.

63. Archibald S, Roy DP, Van Wilgen BW, Scholes RJ. What limits fire? An examination of drivers of burnt area in Southern Africa. Global Change Biology. 2009;15(3):613–630. doi:10.1111/j.1365-2486.2008.01754.x.

64. Mateus D, Gallois JP, Comet JP, Le Gall P. Symbolic modeling of genetic regulatory networks. Journal of Bioinformatics and Computational Biology. 2007;05(02b):627–640. doi:10.1142/S0219720007002850.

65. Pecheur C, Raimondi F. Symbolic Model Checking of Logics with Actions. In: Edelkamp S, Lomuscio A, editors. Model Checking and Artificial Intelligence. Lecture Notes in Computer Science. Berlin, Heidelberg: Springer; 2007. p. 113–128.

66. Alur R, Henzinger TA, Kupferman O. Alternating-time temporal logic. Journal of the ACM. 2002;49(5):672–713. doi:10.1145/585265.585270.

67. Cattelino PJ, Noble IR, Slatyer RO, Kessell SR. Predicting the multiple pathways of plant succession. Environmental Management. 1979;3(1):41–50. doi:10.1007/BF01867067.

68. Pettersson P. Modelling and Verification of Real-Time Systems Using Timed Automata : Theory and Practice [PhD Thesis]. Uppsala University, Department of Computer Systems; 1999.

69. Alur R, Dill DL. A theory of timed automata. Theoretical Computer Science. 1994;126(2):183–235. doi:10.1016/0304-3975(94)90010-8.

70. Cerone A, Maggiolo-Schettini A. Time-based expressivity of time Petri nets for system specification. Theoretical Computer Science. 1999;216(1):1–53. doi:10.1016/S0304-3975(98)00008-5.

71. Bouyer P, Fahrenberg U, Larsen KG, Markey N, Ouaknine J, Worrell J. Model Checking Real-Time Systems. In: Clarke EM, Henzinger TA, Veith H, Bloem R, editors. Handbook of Model Checking. Cham: Springer International Publishing; 2018. p. 1001–1046. Available from: https://doi.org/10.1007/978-3-319-10575-8_29.

72. Alur R, Courcoubetis C, Dill D. Model-Checking in Dense Real-Time. Information and Computation. 1993;104(1):2–34. doi:10.1006/inco.1993.1024.

73. Koymans R. Specifying real-time properties with metric temporal logic. Real-Time Systems. 1990;2(4):255–299. doi:10.1007/BF01995674.

74. Yeakel JD, Pires MM, de Aguiar MAM, O’Donnell JL, Guimarães PR, Gravel D, et al. Diverse interactions and ecosystem engineering can stabilize community assembly. Nature Communications. 2020;11(1):3307. doi:10.1038/s41467-020-17164-x.

75. Baier C, de Alfaro L, Forejt V, Kwiatkowska M. Model Checking Probabilistic Systems. In: Clarke EM, Henzinger TA, Veith H, Bloem R, editors. Handbook of Model Checking. Cham: Springer International Publishing; 2018. p. 963–999. Available from: https://doi.org/10.1007/978-3-319-10575-8_28.

